# A proxy measure of striatal dopamine predicts individual differences in temporal precision

**DOI:** 10.1101/2022.01.21.477273

**Authors:** Renata Sadibolova, Luna Monaldi, Devin B. Terhune

**Author notes:** Address correspondence to: Renata Sadibolova, Department of Psychology, Goldsmiths, University of London, 8 Lewisham Way, London, UK, SE14 6NW. These authors contributed equally to this work (Renata Sadibolova, Luna Monaldi).

## Abstract

The perception of time is characterized by pronounced variability across individuals, with implications for a diverse array of psychological functions. The neurocognitive sources of this variability are poorly understood but accumulating evidence suggests a role for inter-individual differences in striatal dopamine levels. Here we present a pre-registered study that tested the predictions that spontaneous eye blink rates, which provide a proxy measure of striatal dopamine availability, would be associated with aberrant interval timing (lower temporal precision or overestimation bias). Neurotypical adults (*N*=69) underwent resting state eye tracking and completed visual psychophysical interval timing and control tasks. Elevated spontaneous eye blink rates were associated with poorer temporal precision but not with inter-individual differences in perceived duration or performance on the control task. These results signify a role for striatal dopamine in variability in human time perception and can help explain deficient temporal precision in psychiatric populations characterized by elevated dopamine levels.

## Introduction

Human adults display pronounced heterogeneity in interval timing (Wiener et al., 2014), which impacts a diverse array of basic and complex psychological functions from precise subsecond motor control to suprasecond cognitive decision making (Merchant & Georgopoulos, 2006; Sohn & Carlson, 2003). However, the neurocognitive factors underlying timing variability remain poorly understood.

Multiple strands of evidence implicate the striatal dopamine system in interval timing and timing heterogeneity (for reviews, see Agostino & Cheng, 2016; Coull et al., 2011). For example, psychiatric populations characterized by aberrant striatal dopamine profiles exhibit atypical timing (Allman & Meck, 2012). In schizophrenia, which is characterized by a hyper-responsive dopamine system (Howes et al., 2015) and elevated striatal D2-receptor availability (Seeman, 2013), patients reliably display poorer temporal precision (variance of perceived intervals across trials) (Thoenes & Oberfeld, 2017; Ueda et al., 2018). One interpretation of these impairments is that elevated dopamine leads to the over-weighting of priors under uncertainty, which will produce a flattening of psychometric slopes and a corresponding reduction in temporal precision. In support of this proposal, a previous study found that controls and patients with schizophrenia exhibited timing migration effects towards context tone intervals (priors) drawn from low-variance (low uncertainty) distributions (Cassidy et al., 2018). Critically, unlike controls off amphetamine, patients and controls on amphetamine exhibited such timing performance even after the precision of priors was reduced in a high-variance (high uncertainty) context condition.

Conversely, striatal dopamine blockage may diminish signalling of a precise prior. Tomassini et al. (2016, 2019) presented participants with foreperiods between warning and go stimuli, when they were required to respond quickly and accurately. Foreperiod intervals were drawn from distributions with high- or low-means and variances. Critically, low-variance conditions yielded higher predictability of foreperiod offset (prior) and concomitantly improved response times and temporal precision. The administration of haloperidol reduced this advantage, independently of a motor impairment (Tomassini et al., 2016), suggesting that dopamine may mediate the ability to extract temporal expectations (priors) across trials.

Notably, some of these results do not align with the predictions of pacemaker-accumulator models of interval timing (Gibbon et al., 1997; Treisman & Brogan, 1992). In these models, dopamine is hypothesized to affect the speed of a putative internal clock consisting of a pacemaker emitting pulses and an accumulator collecting these pulses. A faster clock (higher dopamine) would be expected to generate more pulses resulting in duration overestimation and finer temporal resolution (superior precision). An abundance of pharmacological and animal research aligns with these predictions (for reviews, see Agostino & Cheng, 2016; Coull, Cheng, et al., 2011) but the evidence is not without controversy. Whereas the D2-receptor agonist quinpirole has been shown to attenuate temporal precision (Santi et al., 2001), another agonist pergolide improved temporal sensitivity (Rammsayer, 2009). Poorer temporal precision was also observed with the dopamine antagonist haloperidol (Coull et al., 2011), but not with sulpiride (Rammsayer, 1997). Moreover, the majority of pharmacological studies do not include appropriate control tasks (Buhusi & Meck, 2002; Maricq & Church, 1983; Rammsayer, 1989a, 1989b, 1997), suggesting that timing deficits may be mediated by non-timing cognitive effects (Coull, 2014; Rammsayer, 1997; Saeedi et al., 2006).

Evidence for a role of dopamine in temporal accuracy (proximity of temporal estimates and stimulus intervals) is similarly mixed. Although some clinical evidence implicates dorsal striatum in temporal accuracy (Allman & Meck, 2012), the recent literature suggests only a weak tendency toward aberrant accuracy in schizophrenia (Thoenes & Oberfeld, 2017) and in Parkinson’s disease (Terao et al., 2021), which are respectively characterized by elevated and diminished dopamine levels. By contrast, pharmacological modulation of dopamine synthesis reliably alters timing accuracy (Coull et al., 2011). For instance, rodents administered dopamine agonists exhibit leftward shifts of psychometric functions, suggesting subjective dilation of time, whereas the administration of antagonists is associated with rightward shifts reflecting temporal contraction (Buhusi & Meck, 2002; Maricq & Church, 1983; Matell et al., 2004). More recently, however, a study using optogenetics reported temporal dilation in response to attenuated midbrain dopamine levels (Soares et al., 2016; see Mikhael & Gershman, 2019, for a proposed reconcillation of these findings).

Free from pharmacological intervention, *baseline* striatal dopamine concentrations (and D2-receptor availability) fluctuate over the course of hours, albeit at much lower magnitude scales (Ferris et al., 2014). Whether the clinical and pharmacological observations extend to the neurotypical baseline is not known. Previous research attributed poorer temporal precision to carriers of a genetic allele associated with reduced density of D2-receptors (Wiener et al., 2011, 2014). However, insofar as the prevalence of this allele is only ∼20-27% in the general population (National Centre for Biotechnology Information, 2021), the relationship between baseline dopamine and temporal perception remains poorly understood.

A common feature of studies of the association between dopamine and interval timing is that they seldom extend to human neurotypical timing free from experimentally induced perturbation of dopaminergic activity, mostly due to the invasive nature of available methods (e.g., Kishida et al., 2016). Yet, such investigations would complement current understanding of timing mechanisms and provide a benchmark for studies with clinical populations. This issue has been addressed in investigations of the role of dopamine in cognition with the use of spontaneous eye blink rates (EBR) as a dopamine proxy (for a review, see Jongkees & Colzato, 2016), given its association with the availability of dorsostriatal D2-receptors (Elsworth et al., 1991; Groman et al., 2014; Karson, 1983; Kleven & Koek, 1996; Taylor et al., 1999; but see also Dang et al., 2017). We previously reported that EBR covaries with intra-individual fluctuations in perceived duration of auditory and visual intervals (Terhune et al., 2016), showing temporal dilation for trials following spontaneous eyeblinks, as would be expected from some pharmacological studies (Buhusi & Meck, 2002; Maricq & Church, 1983; Matell et al., 2004). By contrast, temporal precision did not statistically differ after blinks although we observed a trend for poorer precision in post-blink trials (Terhune et al., 2016).

This pre-registered study sought to expand upon previous observations by investigating how EBR relates to *inter*-individual variability in human neurotypical interval timing. Toward this end, neurotypical adults underwent resting state eye-tracking and completed a visual psychophysical timing task (temporal bisection) and a control task (color bisection) to assess the cognitive specificity of associations between EBR and perception. Drawing on our previous results (Terhune et al., 2016), we expected that EBR, with higher rates reflecting greater striatal dopamine receptor availability (Groman et al., 2014), would be associated with a relative tendency to overestimate stimulus intervals. Terhune et al. (2016) also observed consistent trends for poorer precision in post-blink trials. Based on their observations and the links between elevated dopamine and attenuated temporal precision in schizophrenia (Thoenes & Oberfeld, 2017), we hypothesized that EBR would be associated with poorer temporal precision. An alternative prediction, derived from the striatal beat frequency model (Matell & Meck, 2004), is that EBR would be associated with superior temporal precision.

## Methods

### Participants

75 healthy adults (82.66% female, 17.33% male) between 18 and 45 years old (*M*_Age_=23.07, *SD*=4.22) with at least one year of post-secondary education (*M*=3.43, *SD*=2.21) provided informed written consent to participate in this study in accordance with approval by a local departmental ethics committee. Participants were right-handed, had no history of neurological or psychiatric disorders and had normal or corrected-to-normal vision. Sample size was estimated *a priori* on the basis of a pilot study using G*Power 3.1 (Faul et al., 2009), using the parameters of *r*=.33 (unpublished pilot data), 1-β=.80, α=.05 (two-tailed), which resulted in a required sample size of 69 participants. We pre-specified a target sample size of 75 participants, in order to account for attrition. The study was preregistered prior to data collection on Open Science Framework (https://osf.io/fzdbv).

### Resting state eye-tracking

The detailed eye-tracking protocol has been described elsewhere (Terhune et al., 2016). Participants attended to a fixation cross at the center of a computer monitor whilst their resting state right eye movements were tracked using an EyeLink 1000 (SR Research, Ontario, Canada). Their head movements were minimized using a chinrest. The participant’s gaze location was monitored in real-time by a trained experimenter on a separate monitor out of participant’s view. The data were monocularly sampled with the right eye at a rate of 500Hz. A blink was defined as a period in which a pupil was not detected for three or more consecutive samples.

### Temporal bisection task

Participants completed a visual temporal bisection task that entailed learning two anchor intervals (300 and 967ms) in a training phase. In a subsequent testing phase, participants were presented with colored circles of varying intervals and judged whether they were closer in duration to the trained short or long anchor intervals. Trials consisted of a jittered interstimulus interval (blank screen) drawn from a truncated Poisson distribution (200-500ms), a circle (2.7cm, ∼2° of visual field) that randomly flickered between blue and red at 60Hz and varied in duration (300, 433, 567, 700, 833, or 967ms), a second interstimulus interval (blank screen; 300ms), and a two-alternative forced choice judgment prompt (S L or L S [S=short; L=long]) (see Figure S1 in supplemental materials). There were six different color proportion sets at each stimulus interval, as detailed in *Color bisection task*. Participants responded by pressing one of two keys using their index and middle fingers.

### Color bisection task

A color bisection task was administered to control for attention and working memory demands in the temporal bisection task (Coull, 2014). The stimulus set and trial sequence were identical to those in the temporal bisection task except the training phase, which involved training with two anchor colors (mostly-red: 22% blue and mostly-blue: 78% blue) and the judgment prompt (B R or R B [B=blue, R=red]). At the prompt, participants judged whether the preceding flickering circle was closer to the previously learned blue or red anchor stimuli. Colors were drawn from six proportion sets of blue and red circles (22, 34, 45, 55, 66, or 78% blue). Each color stimulus included equal proportions of the six intervals in the temporal bisection task.

### Procedure

Participants first underwent the resting state eye-tracking, which involved sitting at a desk with their head on a chinrest at ∼55cm distance from the monitor (34×57cm) and ∼40cm distance from the eye tracker camera. They first completed a 9-point calibration procedure and subsequently attended to a centrally presented fixation cross on the monitor for eight minutes, during which their eye movements and pupil diameter were continuously recorded. The experimenter monitored the participant’s eye movements on an external monitor outside of the participant’s view. The room was kept dark during the eye-tracking recording and subsequent tasks.

Participants next completed the temporal and color bisection tasks with stimulus presentation implemented with Psychtoolbox (Brainard, 1997) in MATLAB v. 2018b (MathWorks, Natick, USA). Task order and response key mappings (S L vs. L S and B R vs. R B) were both counterbalanced across participants. The training phases consisted of 20 trials, comprised of equal proportions of each anchor stimulus. Additional trials automatically followed if accuracy was below 80% until the 80% target was reached in the lattermost 20 trials or until the maximum training time of six minutes had passed. In the temporal bisection task, seven participants required more than 20 anchor trials in the training phase and four of them eventually achieved the performance target. In the color bisection task, four participants required additional anchor trials in the training phase, and two of them subsequently reached the performance target. The five participants who did not reach the performance target completed both training phases with >70% accuracy.

Participants subsequently completed 3 blocks of 76 trials of each task. Experimental blocks began with two reminders of each anchor stimulus (4 trials); the remaining 72 trials included 16 repetitions of the 4 middle stimuli and 4 repetitions of each anchor stimulus. In both tasks, participants were instructed to focus on the center of the monitor and only use the interval or color information to guide their judgment and to avoid other strategies such as counting or humming. Participants subsequently completed two psychometric measures (hallucination-proneness and frequency of drug use) that will be reported independently.

### Analyses

All analyses were performed in MATLAB. Eye-tracking data were used to compute the mean number of eye blinks per minute (eye blink rate; EBR). The EBR includes only the last 5 minutes of the recording as the first 3 minutes were discarded to account for adaptation to the room lighting. Task performance was assessed by computing the proportion of “long” [*p*(long)] and “red” [*p*(red)] responses at each stimulus level in the temporal and color bisection tasks, respectively. We fitted logistic functions to these values in individual participants using maximum likelihood estimation as implemented in the Palamedes toolbox (Kingdom & Prins, 2016) in order to estimate the alpha (bisection point [BP]) and beta (slope) parameters of the psychometric function (guess rates and lapse rates were set at 0). The fit of psychometric functions was assessed by computing a *p*-value for deviance (*p*Dev), based on 1,000 permutations. In each task, we estimated the bisection point (BP), which corresponds to the estimated stimulus level that is perceived as equidistant to the two anchor stimuli (lower values reflect relative overestimation of intervals [temporal bisection] and relative overestimation of redness [color bisection]). We additionally estimated the Weber fraction (WF), which is the difference limen proportional to the BP, with the former given as half of the difference between the intervals (or color proportions) corresponding to 75% and 25% of the *p*(long) (or *p*(red)) response proportions on the fitted psychometric function (lower values reflect superior precision).

Two participants’ eye-tracking data were not recorded due to technical errors involving the eye-tracker. We additionally excluded 4 participants in the temporal bisection task and 6 participants in the color bisection task due to poor fit of the psychometric functions, *p*Dev<.05. These exclusions resulted in final sample sizes of 69 and 67 for the temporal and color bisection tasks, respectively. EBRs were correlated with psychophysical parameters using the Robust Correlation toolbox in MATLAB (Pernet et al., 2013). Spearman correlations (*r*_s_) and partial Spearman correlations (*r*_ps_) were used throughout due to violations of distribution normality or homoscedasticity. We report correlations pre- and post-multivariate outlier removal for completeness (the latter are reported in Supplemental materials, section S6. Bivariate outliers). Correlations are supplemented with bias-corrected and accelerated confidence intervals using Bootstrapping (10,000 samples) (Efron, 1987). Correlations were compared across tasks by standardizing coefficients (Fisher z-transform; Myers & Sirois, 2014) and computing the Bootstrap 95% confidence intervals of the correlation coefficient difference, with intervals non-overlapping with zero denoting significance.

For each correlation, we additionally report the Bayes factor (BF_10_) quantifying the likelihood of the data under the experimental hypothesis H_1_ relative to the likelihood of the data under the null hypothesis H_0_. The BF_+0_ and BF_-0_ denote the hypothesized positive and negative correlations, respectively. Conventionally, BF values greater than 3 and less than 1/3 denote moderate or greater evidence for H_1_ and H_0,_ respectively, whereas intermediate values indicate an ambiguous result (Wagenmakers et al., 2016). BFs were computed in JASP (JASP Team, 2019) with pre-specified default parameters of the H_1_ and H_0_ distributions (Wagenmakers et al., 2016) and their robustness was verified using a BF sensitivity analysis with different H_1_ priors (section S2 in supplemental materials). When different priors yielded markedly different BF results, the ambiguity of the evidence was acknowledged.

## Results

Our primary analyses tested the predictions that EBR would be selectively associated with performance in the temporal bisection task but not in the control (color bisection) task. In particular, we expected that higher EBR (reflecting elevated striatal dopamine receptor availability; Jongkees & Colzato, 2016) would be associated with higher WFs (reflecting poorer temporal precision) and shorter BPs (reflecting longer perceived duration) (see Figure 1 for summary information of the data).

**Figure 1.**
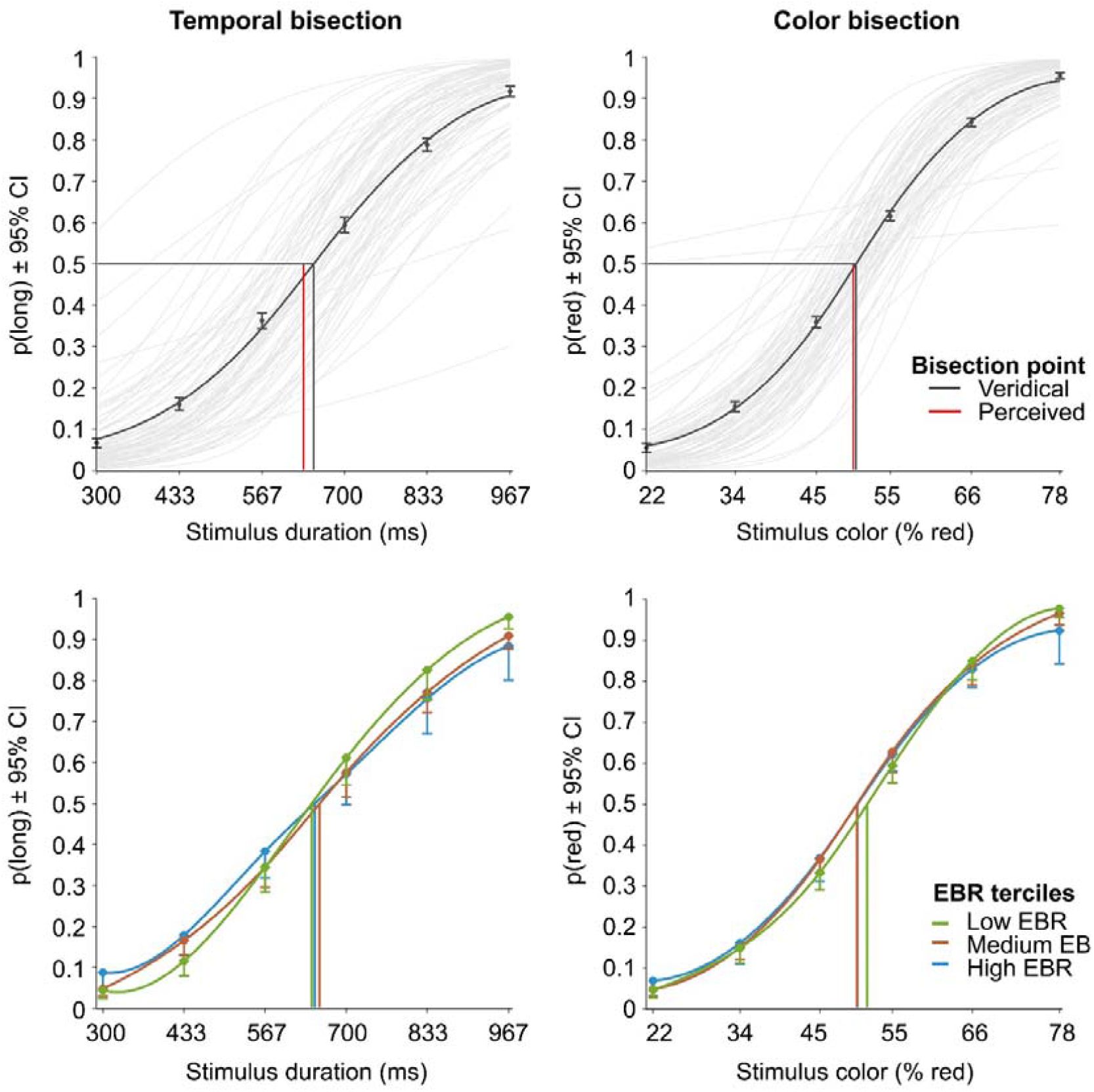
PF models of task performance. The lines in saturated colors show spline-interpolated sample means of actual p(response) for each stimulus. **Top row**: The pale grey lines represent individual PFs for all participants and the vertical lines denote veridical and perceived BPs. Circular markers and error bars denote sample means and 95% bootstrapped confidence intervals for the proportions of long [p(long)] or red [p(red)] responses. **Bottom row:** Group PFs for low, medium, and high EBR terciles. To improve legibility of the overlapping error bars, only the lower limits are plotted.

### EBR and temporal precision

In support of our first prediction, EBR positively correlated with WFs in the temporal bisection task, *r*_s_=.28 [Bootstrap 95% CI: .04, .49], *p*=.019 (*N*=69) (see Figure 2). This correlation is similarly reflected in Figure 1 (bottom left), where the first tercile, comprising participants with the lowest EBR, displayed the steepest slopes, reflecting superior temporal precision. The BF_+0_ was 4.02, suggesting that these data were four times more likely to be observed under our hypothesis than the null hypothesis. By contrast, EBR did not significantly correlate with WFs in the control (color bisection) task, *r*_s_=.08 [-.17, .32], *p*=.50 (*N*=67), with Bayesian evidence in favor of the null hypothesis, BF_+0_=.31. The robustness of these results is supported by BF sensitivity analyses and the strength of evidence further increased after excluding bivariate outliers (see Supplemental Materials). Finally, the temporal specificity of this effect was corroborated by the correlation between temporal WFs and EBR being significantly greater than that between color WFs and EBRs, *r*_s(diff)_=.33 [Bootstrap 95% CI: .05, .54], *t*(62)=2.52, *p*=.010, *d*_z_=.32.

**Figure 2.**
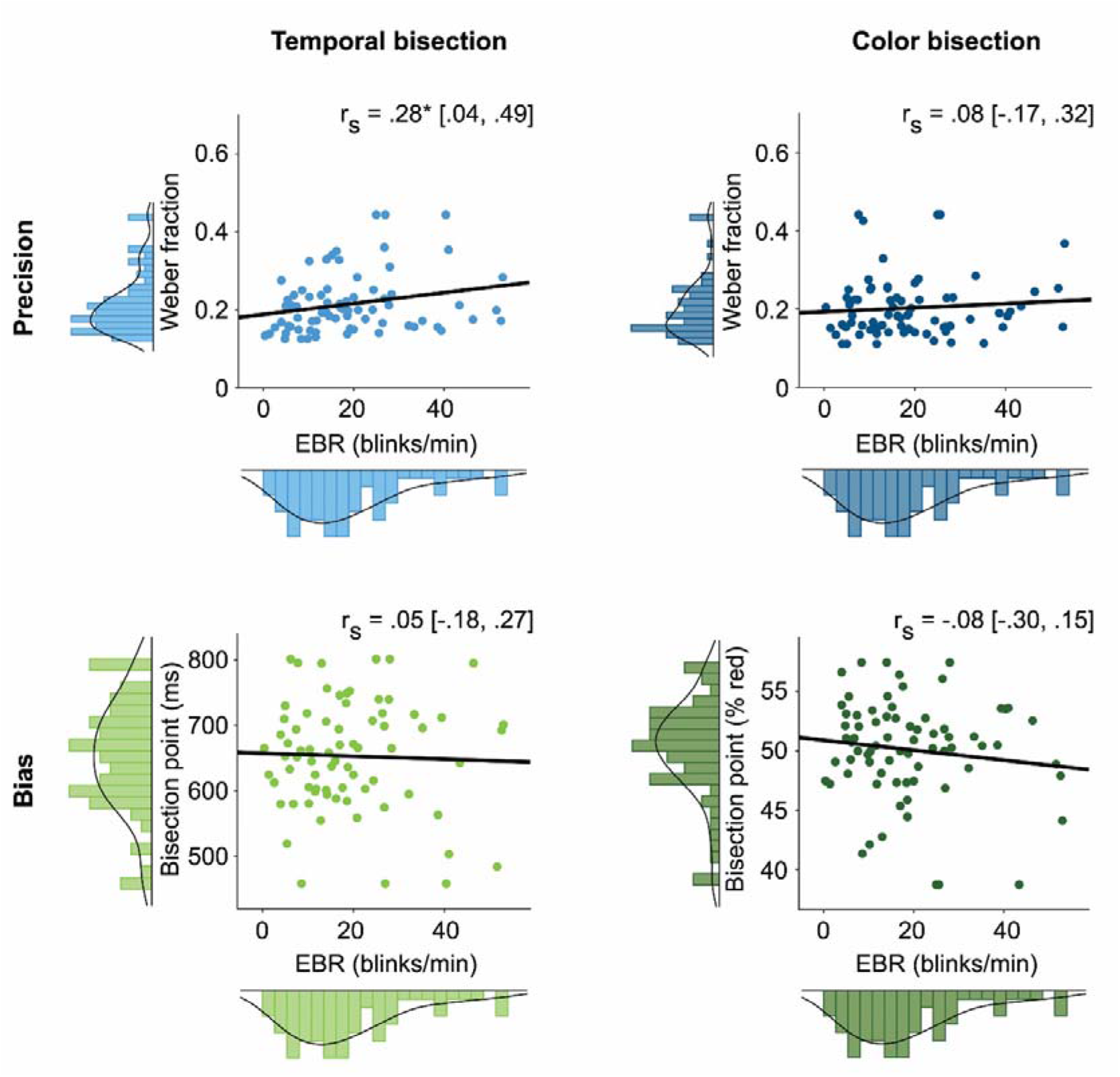
Individual Weber fractions (WFs; blue) and Bisection points (BPs; green) as a function of spontaneous eye blink rate (EBR). Square brackets denote bootstrapped 95% confidence intervals. Plots show original non-ranked data with winsorized outliers, and the least-squares line fit for visualization purposes. *r*_s_ = Spearman correlation.

A series of unregistered correlational analyses were conducted to exclude alternative interpretations of the reported effects. Temporal and color WFs were positively correlated, *r*_s_=.31 [.09, .51], *p*=.011 (*N*=65), BF_+0_=6.77. A semi-partial Spearman correlation between temporal WFs and EBR (after partialling out shared variance between temporal and color WFs), was significant, *r*_ps_ =.33 [.05, .53], *p*=.009 (*N*=63), BF_+0_=9.62. The association between temporal WFs and EBR similarly remained stable when partialling out the shared variance between BPs and WFs (see Supplemental Materials). Conversely, the association between EBR and color WFs remained non-significant after controlling for temporal WFs, *r*_ps_ =-.11 [-.35, .13], *p*=.39 (*N*=63), BF_+0_=.09. Further, since larger WFs are associated with flatter psychometric function slopes and as such reflect increased task difficulty, we compared the WFs across tasks. We observed a non-significant difference with evidence for the null hypothesis, *t*(74)=1.21, *p*=.23, *d*_z_=.14, BF_10_=.26, suggesting that the tasks were appropriately matched in difficulty.

Cumulatively, these results suggest that although temporal and color precision were associated, EBR is selectively associated with poorer temporal precision, but not with color precision.

### EBR and temporal bias

In contrast with our prediction, EBR did not significantly correlate with temporal BPs (Figure 2), *r*_s_=.03 [-.23, .29], *p*=.78 (*N*= 69), with clear evidence in favor of the null hypothesis, BF_-0_=.14. The same held for color BPs, *r*_s_=-.12 [-.36, .13], *p*=.31 (*N*=67), albeit with more ambiguous Bayesian evidence, BF_-0_=.40. The two correlations did not significantly differ, *r*_s(diff)_=.04 [Bootstrap 95% CI: -.31, .38], *t*(62)=.34, *p*=.74, *d*_z_=.04.

BPs in the two tasks did not significantly correlate *r*_s_=.10 [-.17, .35], *p*=.43 (*N*=65), with evidence for the null hypothesis, BF_10_=.22. Controlling for the influence of the other task did not change the association between EBR and temporal BPs, *r*_ps_=-.03 [- .28, .23], *p*=.80 (*N*=63), BF_-0_=.24, or between EBR and color BPs, *r*_ps_=-.09 [-.33, .15], *p*=.49 (*N*=63), BF_-0_=.32. Taken together, these results suggest that EBR is unrelated to inter-individual variation in perceived duration.

### EBR and response times

Although the color task allowed for dissociating non-specific memory effects, the decisional components of task performance were further modelled using hierarchical drift diffusion modelling (DDM; Ratcliff & McKoon, 2008; Wiecki et al., 2013). DDM was fitted to responses and response times to decompose the data into parameters reflecting the decision process (prior bias, speed of evidence accumulation, decision thresholds and non-decisional perceptual and motor processes). We assessed the association between EBR and these parameters and we observed non-significant results (see Supplemental Materials).

## Discussion

This study was motivated by the wealth of pharmacological and clinical data linking interval timing to striatal dopamine (Buhusi & Meck, 2002; Rammsayer, 1993; Thoenes & Oberfeld, 2017) but a corresponding dearth of evidence regarding the role of dopamine in baseline neurotypical temporal cognition. We found that elevated striatal D2-receptor availability, inferred non-invasively from spontaneous eye blink rates (EBR), was selectively associated with poorer temporal precision, but not overestimation or precision or bias in a difficulty- and stimulus-matched control task. Moreover, although temporal and color bisection tasks tax partially overlapping cognitive processes (Coull, 2014), the association between temporal precision and EBR was independent of color precision. These results expand upon research implicating dopamine systems in interval timing (Coull et al., 2011) by demonstrating a similar link without pharmacological intervention in healthy individuals.

Higher dopamine was previously suggested to drive increased reliance on expectations (priors) in interval timing (Cassidy et al., 2018). Cassidy et al. (2018) manipulated reliance on temporal priors in a temporal reproduction task in patients with schizophrenia and controls. They replicated the standard migration effect across groups (Jazayeri & Shadlen, 2010) and observed that timing performance in patients and controls on amphetamine was influenced even when priors became less reliable (Cassidy et al., 2018). On the basis of these results, we suggest that higher EBR in the present study was associated with overreliance on priors, particularly under uncertainty (stimulus intervals close to the middle interval). This led responses to drift closer to the prior (mean of the interval range), resulting in lower precision and higher WFs. This result appears to diverge from Terhune et al.’s (2016) observation of *no* significant association between trial-by-trial blink patterns and temporal *precision*. However, that study did observe a trend in the direction of poorer precision in post-blink trials, which aligns with the present results.

This interpretation aligns with results demonstrating that schizophrenia is characterized by reduced temporal precision (Thoenes & Oberfeld, 2017), and elevated dopamine (Howes et al., 2015), D2-receptor availability (Seeman, 2013), and EBR (Adamson, 1995), as well as reduced temporal precision in its subclinical expression (schizotypy; Ferri et al., 2017). However, whilst models of schizophrenia acknowledge some role of general motivational processes, we show an effect specific to timing, as reflected by a significantly lower association between EBR and color precision. Further, we replicated our central result after adjusting for shared variance in precision in the two tasks.

An alternative interpretation of our results in line with the striatal beat-frequency model of timing (SBF; Matell & Meck, 2004) is that higher dopamine impairs coincidence detection by striatal spiny neurons (Buhusi & Meck, 2005; Urakubo et al., 2020), which would lead to a reduced precision regarding the interval onset or offset times (Paton & Buonomano, 2018). Whereas these conclusions are speculative, there are some indications supporting this possibility. Allman and Meck (2012) suggested that atypical stimulus onsets and cortical asynchronization may explain timing variability in schizophrenia, reflecting disturbed coincidence detection and starting (or ‘resetting’) of the striatal interval clock under the SBF model.

Insofar as the speed of a putative clock is hypothesized to increase with elevated D2-receptor activity (MacDonald & Meck, 2005), our results are inconsistent with this model’s prediction that a higher clock speed, mediated by elevated dopamine, relates to improved temporal precision. The SBF model proposes that activation of striatal neurons reflects their sensitivity to a pattern of glutamatergic corticostriatal signals (MacDonald & Meck, 2005). There is evidence that dopamine, and D2-receptors specifically, play a role in the filtering of more active glutamatergic corticostriatal inputs (Bamford et al., 2004). Conversely, interval training has been shown to ‘blunt’ the effects of dopaminergic drugs and this ‘dopamine-insensitive’ state was reversed by administration of ketamine, an NMDA (glutamate) receptor antagonist (Cheng et al., 2007). Importantly, Cheng et al. (2007) highlighted that the transition to a dopamine-insensitive state was similar to general observations of striatal neurons becoming silent once a reward becomes predictable with training (Schultz, 1998). Similarly, Wang et al. (2020) recently suggested that the noise in corticostriatal circuits that underlies timing variability is subject to adjustments through reinforcement learning. This overlap between learning and timing may help to reconcile our seemingly discrepant results.

At first glance, our results demonstrating no association between temporal bias and EBR seem at odds with the previously reported association between trial-by-trial eyeblinks and the tendency to overestimate stimulus intervals (Terhune et al., 2016). Our results suggest that individual differences in striatal dopamine do not contribute to inter-individual variability in temporal bias, which is consistent with the available evidence from clinical populations characterized by dopamine dysregulation (Terao et al., 2021; Thoenes & Oberfeld, 2017; but see Ueda et al., 2018). Alternatively, given that the link between temporal accuracy and dopamine (D2) expression has typically been studied in conditions when dopamine availability markedly deviated from a neurotypical baseline, our data may demonstrate the null effect across much smaller inter-individual variations at baseline.

Some recent developments suggest that interval timing may not require internally-driven mechanisms involving basal ganglia and that exteroceptive perceptual content alone may be the principal determinant of subjective time (Roseboom et al., 2019; Suárez-Pinilla et al., 2019). This model would not strictly preclude dopamine-mediated timing; it would, however, favor a more complex and indirect interpretation, for instance via dopamine-affected signaling within local circuits and global networks that impacts attention but also timing specifically (Coull et al., 2012; Nagano-Saito et al., 2008; Shafiei et al., 2019). Nevertheless, whereas sparse population coding, with neuronal populations activating in sequence over the course of the timed interval, was reported in regions such as orbitofrontal cortex and secondary motor cortex (Bakhurin et al., 2017; Zhou et al., 2020), *striatal* activity exhibits the highest degree of sequentiality for the parsing of intervals by biologically constrained decoder networks and therefore provides the most optimized set of signals to readout duration (Zhou et al., 2020).

EBR shows considerable promise as a non-invasive proxy of striatal dopamine in timing research if its limitations are acknowledged. Although the evidence shows it to be a viable proxy for D2-receptor availability (Groman et al., 2014; Jongkees & Colzato, 2016), it is nonetheless only an indirect measure and the strength of the association between EBR and D2-receptor availability will therefore modulate our findings. It remains poorly understood how stable this association is across the typical range as well as for more extreme high and low dopamine levels in human striatum as well as how stable it is across individuals and over time. Although we only recorded EBR at baseline prior to the completion of the perceptual tasks, previous research shows that EBR is stable over short time periods (>1 hour) (Barbato et al., 2000). Since Terhune et al. (2016) observed their effects both in sub- and suprasecond interval ranges, future studies should assess whether the current findings extend to suprasecond timing. Finally, our use of basic visual stimuli does not allow for an assessment of perceptual context in the shaping of subjective timing in a manner that is enabled with the use of more ecologically valid stimuli (Roseboom et al., 2019; van Rijn, 2018).

To summarize, the association between EBR and interval timing was timing- and precision-specific and thereby builds on research implicating dopamine in interval timing (Coull et al., 2011) and extends this to individual differences in the neurotypical population. Altogether, our results complement studies demonstrating associations between EBR and cognitive-perceptual functions subserved by dopamine systems (Jongkees & Colzato, 2016), and they attest to the utility of EBR as a proxy measure of dopamine in time perception research.

## Declarations

### Data availability

The data and materials are available at https://osf.io/jxc3f.

### Code availability

Custom scripts are available at https://osf.io/jxc3f.

### Authors’ contributions

All authors designed the research, LM collected the data, RS analyzed the data, RS and DBT drafted the manuscript, and all authors approved the final manuscript for submission.

## Acknowledgments

This paper is dedicated to the memory of Warren H. Meck.

## References

Adamson, T. A. (1995). Changes in blink rates of Nigerian schizophrenics treated with chlorpromazine. West African Journal of Medicine, 14(4), 194–197.

Agostino, P. V., & Cheng, R.-K. K. (2016). Contributions of dopaminergic signaling to timing accuracy and precision. Current Opinion in Behavioral Sciences, 8, 153–160. https://doi.org/10.1016/j.cobeha.2016.02.013

Allman, M. J., & Meck, W. H. (2012). Pathophysiological distortions in time perception and timed performance. Brain, 135(3), 656–677. https://doi.org/10.1093/brain/awr210

Bakhurin, K. I., Goudar, V., Shobe, J. L., Claar, L. D., Buonomano, D. V., & Masmanidis, S. C. (2017). Differential encoding of time by prefrontal and striatal network dynamics. Journal of Neuroscience, 37(4), 854–870. https://doi.org/10.1523/JNEUROSCI.1789-16.2016

Bamford, N. S., Zhang, H., Schmitz, Y., Wu, N. P., Cepeda, C., Levine, M. S., Schmauss, C., Zakharenko, S. S., Zablow, L., & Sulzer, D. (2004). Heterosynaptic dopamine neurotransmission selects sets of corticostriatal terminals. Neuron, 42(4), 653–663. https://doi.org/10.1016/S0896-6273(04)00265-X

Barbato, G., Ficca, G., Muscettola, G., Fichele, M., Beatrice, M., & Rinaldi, F. (2000). Diurnal variation in spontaneous eye-blink rate. Psychiatry Research, 93(2), 145–151. https://doi.org/10.1016/S0165-1781(00)00108-6

Brainard, D. H. (1997). The Psychophysics Toolbox. Spatial Vision, 10(4), 433–436. https://doi.org/10.1163/156856897X00357

Buhusi, C. V., & Meck, W. H. (2002). Differential effects of methamphetamine and haloperidol on the control of an internal clock. Behavioral Neuroscience, 116(2), 291–297. https://doi.org/10.1037/0735-7044.116.2.291

Buhusi, C. V., & Meck, W. H. (2005). What makes us tick? Functional and neural mechanisms of interval timing. Nature Reviews Neuroscience, 6(10), 755–765. https://doi.org/10.1038/nrn1764

Cassidy, C. M., Balsam, P. D., Weinstein, J. J., Rosengard, R. J., Slifstein, M., Daw, N. D., Abi-Dargham, A., & Horga, G. (2018). A perceptual inference mechanism for hallucinations linked to striatal dopamine. Current Biology, 28(4), 503-514.e4. https://doi.org/10.1016/j.cub.2017.12.059

Cheng, R.-K., Ali, Y. M., & Meck, W. H. (2007). Ketamine “unlocks” the reduced clock-speed effects of cocaine following extended training: Evidence for dopamine–glutamate interactions in timing and time perception. Neurobiology of Learning and Memory, 88(2), 149–159. https://doi.org/10.1016/j.nlm.2007.04.005

Coull, J. T. (2014). Getting the Timing Right: Experimental Protocols for Investigating Time with Functional Neuroimaging and Psychopharmacology. In H. Merchant & V. de Lafuente (Eds.), Advances in Experimental Medicine and Biology (pp. 237–264). Springer Science+Business Media. https://doi.org/10.1007/978-1-4939-1782-2_13

Coull, J. T., Cheng, R.-K., & Meck, W. H. (2011). Neuroanatomical and neurochemical substrates of timing. Neuropsychopharmacology, 36(1), 3–25. https://doi.org/10.1038/npp.2010.113

Coull, J. T., Hwang, H. J., Leyton, M., & Dagher, A. (2012). Dopamine Precursor Depletion Impairs Timing in Healthy Volunteers by Attenuating Activity in Putamen and Supplementary Motor Area. Journal of Neuroscience, 32(47), 16704–16715. https://doi.org/10.1523/JNEUROSCI.1258-12.2012

Dang, L. C., Samanez-Larkin, G. R., Castrellon, J. J., Perkins, S. F., Cowan, R. L., Newhouse, P. A., & Zald, D. H. (2017). Spontaneous Eye Blink Rate (EBR) Is Uncorrelated with Dopamine D2 Receptor Availability and Unmodulated by Dopamine Agonism in Healthy Adults. Eneuro, 4(5), ENEURO.0211-17.2017. https://doi.org/10.1523/ENEURO.0211-17.2017

Efron, B. (1987). Better Bootstrap Confidence Intervals. Journal of the American Statistical Association, 82(397), 171–185. https://doi.org/10.1080/01621459.1987.10478410

Elsworth, J. D., Lawrence, M. S., Roth, R. H., Taylor, J. R., Mailman, R. B., Nichols, D. E., Lewis, M. H., & Redmond, D. E. (1991). D1 and D2 dopamine receptors independently regulate spontaneous blink rate in the vervet monkey. Journal of Pharmacology and Experimental Therapeutics, 259(2), 595–600.

Faul, F., Erdfelder, E., Buchner, A., & Lang, A. (2009). Statistical power analyses using G*Power 3.1: Tests for correlation and regression analyses. Behavior Research Methods, 41(4), 1149–1160. https://doi.org/10.3758/BRM.41.4.1149

Ferri, F., Nikolova, Y. S., Perrucci, M. G., Costantini, M., Ferretti, A., Gatta, V., Huang, Z., Edden, R. A. E., Yue, Q., D’Aurora, M., Sibille, E., Stuppia, L., Romani, G. L., & Northoff, G. (2017). A Neural “Tuning Curve” for Multisensory Experience and Cognitive-Perceptual Schizotypy. Schizophrenia Bulletin, 43(4), 801–813. https://doi.org/10.1093/schbul/sbw174

Ferris, M. J., España, R. A., Locke, J. L., Konstantopoulos, J. K., Rose, J. H., Chen, R., & Jones, S. R. (2014). Dopamine transporters govern diurnal variation in extracellular dopamine tone. Proceedings of the National Academy of Sciences of the United States of America, 111(26), E2751–E2759. https://doi.org/10.1073/pnas.1407935111

Gibbon, J., Malapani, C., Dale, C. L., & Gallistel, C. R. (1997). Toward a neurobiology of temporal cognition: Advances and challenges. Current Opinion in Neurobiology. https://doi.org/10.1016/S0959-4388(97)80005-0

Groman, S. M., James, A. S., Seu, E., Tran, S., Clark, T. A., Harpster, S. N., Crawford, M., Burtner, J. L., Feiler, K., Roth, R. H., Elsworth, J. D., London, E. D., & Jentsch, J. D. (2014). In the Blink of an Eye: Relating Positive-Feedback Sensitivity to Striatal Dopamine D2-Like Receptors through Blink Rate. Journal of Neuroscience, 34(43), 14443–14454. https://doi.org/10.1523/JNEUROSCI.3037-14.2014

Howes, O., McCutcheon, R., & Stone, J. (2015). Glutamate and dopamine in schizophrenia: An update for the 21 st century. Journal of Psychopharmacology, 29(2), 97–115. https://doi.org/10.1177/0269881114563634

JASP Team. (2019). JASP. In [Computer software].

Jazayeri, M., & Shadlen, M. N. (2010). Temporal context calibrates interval timing. Nature Neuroscience, 13(8), 1020–1026. https://doi.org/10.1038/nn.2590

Jongkees, B. J., & Colzato, L. S. (2016). Spontaneous eye blink rate as predictor of dopamine-related cognitive function—A review. Neuroscience & Biobehavioral Reviews, 71, 58–82. https://doi.org/10.1016/j.neubiorev.2016.08.020

Karson, C. N. (1983). Spontaneous Eye-blink Rates and Dopaminergic Systems. Brain, 106(3), 643–653. https://doi.org/10.1093/brain/106.3.643

Kingdom, F. A. A., & Prins, N. (2016). Psychophysics. In Psychophysics: A Practical Introduction: Second Edition. Elsevier. https://doi.org/10.1016/C2012-0-01278-1

Kishida, K. T., Saez, I., Lohrenz, T., Witcher, M. R., Laxton, A. W., Tatter, S. B., White, J. P., Ellis, T. L., Phillips, P. E. M., & Montague, P. R. (2016). Subsecond dopamine fluctuations in human striatum encode superposed error signals about actual and counterfactual reward. Proceedings of the National Academy of Sciences, 113(1), 200–205. https://doi.org/10.1073/pnas.1513619112

Kleven, M. S., & Koek, W. (1996). Differential effects of direct and indirect dopamine agonists on eye blink rate in cynomolgus monkeys. Journal of Pharmacology and Experimental Therapeutics, 279(3).

MacDonald, C. J., & Meck, W. H. (2005). Differential effects of clozapine and haloperidol on interval timing in the supraseconds range. Psychopharmacology, 182(2), 232–244. https://doi.org/10.1007/s00213-005-0074-8

Maricq, A. V., & Church, R. M. (1983). The differential effects of haloperidol and methamphetamine on time estimation in the rat. Psychopharmacology, 79(1), 10–15. https://doi.org/10.1007/BF00433008

Matell, M. S., King, G. R., & Meck, W. H. (2004). Differential Modulation of Clock Speed by the Administration of Intermittent Versus Continuous Cocaine. Behavioral Neuroscience, 118(1), 150–156. https://doi.org/10.1037/0735-7044.118.1.150

Matell, M. S., & Meck, W. H. (2004). Cortico-striatal circuits and interval timing: coincidence detection of oscillatory processes. Cognitive Brain Research, 21(2), 139–170. https://doi.org/10.1016/j.cogbrainres.2004.06.012

Merchant, H., & Georgopoulos, A. P. (2006). Neurophysiology of Perceptual and Motor Aspects of Interception. Journal of Neurophysiology, 95(1), 1–13. https://doi.org/10.1152/jn.00422.2005

Mikhael, J. G., & Gershman, S. J. (2019). Adapting the flow of time with dopamine. Journal of Neurophysiology, 121(5), 1748–1760. https://doi.org/10.1152/jn.00817.2018

Myers, L., & Sirois, M. J. (2014). Spearman Correlation Coefficients, Differences between. In Wiley StatsRef: Statistics Reference Online. John Wiley & Sons, Ltd. https://doi.org/10.1002/9781118445112.stat02802

Nagano-Saito, A., Leyton, M., Monchi, O., Goldberg, Y. K., He, Y., & Dagher, A. (2008). Dopamine depletion impairs frontostriatal functional connectivity during a set-shifting task. Journal of Neuroscience, 28(14), 3697–3706. https://doi.org/10.1523/JNEUROSCI.3921-07.2008

National Centre for Biotechnology Information. (2021). rs1800497 RefSNP Report - dbSNP - NCBI. https://www.ncbi.nlm.nih.gov/snp/rs1800497?vertical_tab=true

Paton, J. J., & Buonomano, D. V. (2018). The Neural Basis of Timing: Distributed Mechanisms for Diverse Functions. Neuron, 98(4), 687–705. https://doi.org/10.1016/j.neuron.2018.03.045

Pernet, C. R., Wilcox, R., & Rousselet, G. A. (2013). Robust Correlation Analyses: False Positive and Power Validation Using a New Open Source Matlab Toolbox. Frontiers in Psychology, 3. https://doi.org/10.3389/fpsyg.2012.00606

Rammsayer, T. (1989a). Is There a Common Dopaminergic Basis of Time Perception and Reaction Time? Neuropsychobiology, 21(1), 37–42. https://doi.org/10.1159/000118549

Rammsayer, T. (1989b). Dopaminergic and Serotoninergic Influence on Duration Discrimination and Vigilance. Pharmacopsychiatry, 22(S1), 39–43. https://doi.org/10.1055/s-2007-1014623

Rammsayer, T. (1997). Are There Dissociable Roles of the Mesostriatal and Mesolimbocortical Dopamine Systems on Temporal Information Processing in Humans? Neuropsychobiology, 35(1), 36–45. https://doi.org/10.1159/000119328

Rammsayer, T. (2009). Effects of Pharmacologically Induced Dopamine-Receptor Stimulation on Human Temporal Information Processing. NeuroQuantology, 7(1), 103–113. https://doi.org/10.14704/nq.2009.7.1.212

Rammsayer, T. H. (1993). On dopaminergic modulation of temporal information processing. Biological Psychology, 36(3), 209–222. https://doi.org/10.1016/0301-0511(93)90018-4

Ratcliff, R., & McKoon, G. (2008). The Diffusion Decision Model: Theory and Data for Two-Choice Decision Tasks. Neural Computation, 20(4), 873–922. https://doi.org/10.1162/neco.2008.12-06-420

Roseboom, W., Fountas, Z., Nikiforou, K., Bhowmik, D., Shanahan, M., & Seth, A. K. (2019). Activity in perceptual classification networks as a basis for human subjective time perception. Nature Communications, 10(1), 267. https://doi.org/10.1038/s41467-018-08194-7

Saeedi, H., Remington, G., & Christensen, B. K. (2006). Impact of haloperidol, a dopamine D2 antagonist, on cognition and mood. Schizophrenia Research, 85(1–3), 222–231. https://doi.org/10.1016/j.schres.2006.03.033

Santi, A., Coppa, R., & Ross, L. (2001). Effects of the dopamine D2 agonist, quinpirole, on time and number processing in rats. Pharmacology Biochemistry and Behavior, 68(1), 147–155. https://doi.org/10.1016/S0091-3057(00)00452-4

Schultz, W. (1998). Predictive reward signal of dopamine neurons. In Journal of Neurophysiology (Vol. 80, Issue 1, pp. 1–27). J Neurophysiol. https://doi.org/10.1152/jn.1998.80.1.1

Seeman, P. (2013). Schizophrenia and dopamine receptors. European Neuropsychopharmacology, 23(9), 999–1009. https://doi.org/10.1016/j.euroneuro.2013.06.005

Shafiei, G., Zeighami, Y., Clark, C. A., Coull, J. T., Nagano-Saito, A., Leyton, M., Dagher, A., & Mišić, B. (2019). Dopamine Signaling Modulates the Stability and Integration of Intrinsic Brain Networks. Cerebral Cortex, 29(1), 397–409. https://doi.org/10.1093/cercor/bhy264

Soares, S., Atallah, B. V., & Paton, J. J. (2016). Midbrain dopamine neurons control judgment of time. Science, 354(6317), 1273–1277. https://doi.org/10.1126/science.aah5234

Sohn, M. H., & Carlson, R. A. (2003). Implicit temporal tuning of working memory strategy during cognitive skill acquisition. American Journal of Psychology. https://doi.org/10.2307/1423579

Suárez-Pinilla, M., Nikiforou, K., Fountas, Z., Seth, A. K., & Roseboom, W. (2019). Perceptual Content, Not Physiological Signals, Determines Perceived Duration When Viewing Dynamic, Natural Scenes. Collabra: Psychology, 5(1). https://doi.org/10.1525/collabra.234

Taylor, J. R., Elsworth, J. D., Lawrence, M. S., Sladek, J. R., Roth, R. H., & Redmond, D. E. (1999). Spontaneous Blink Rates Correlate with Dopamine Levels in the Caudate Nucleus of MPTP-Treated Monkeys. Experimental Neurology, 158(1), 214–220. https://doi.org/10.1006/exnr.1999.7093

Terao, Y., Honma, M., Asahara, Y., Tokushige, S., Furubayashi, T., Miyazaki, T., Inomata-Terada, S., Uchibori, A., Miyagawa, S., Ichikawa, Y., Chiba, A., Ugawa, Y., & Suzuki, M. (2021). Time Distortion in Parkinsonism. Frontiers in Neuroscience, 15, 648814. https://doi.org/10.3389/fnins.2021.648814

Terhune, D. B., Sullivan, J. G., & Simola, J. M. (2016). Time dilates after spontaneous blinking. Current Biology, 26(11), R459–R460. https://doi.org/10.1016/j.cub.2016.04.010

Thoenes, S., & Oberfeld, D. (2017). Meta-analysis of time perception and temporal processing in schizophrenia: Differential effects on precision and accuracy. Clinical Psychology Review, 54, 44–64. https://doi.org/10.1016/j.cpr.2017.03.007

Tomassini, A., Pollak, T. A., Edwards, M. J., & Bestmann, S. (2019). Learning from the past and expecting the future in Parkinsonism: Dopaminergic influence on predictions about the timing of future events. Neuropsychologia, 127, 9–18. https://doi.org/10.1016/j.neuropsychologia.2019.02.003

Tomassini, A., Ruge, D., Galea, J. M., Penny, W., & Bestmann, S. (2016). The role of dopamine in temporal uncertainty. Journal of Cognitive Neuroscience, 28(1), 96–110. https://doi.org/10.1162/jocn_a_00880

Treisman, M., & Brogan, D. (1992). Time Perception and the Internal Clock: Effects of Visual Flicker on the Temporal Oscillator. European Journal of Cognitive Psychology, 4(1), 41–70. https://doi.org/10.1080/09541449208406242

Ueda, N., Maruo, K., & Sumiyoshi, T. (2018). Positive symptoms and time perception in schizophrenia: A meta-analysis. Schizophrenia Research: Cognition, 13, 3–6. https://doi.org/10.1016/j.scog.2018.07.002

Urakubo, H., Yagishita, S., Kasai, H., & Ishii, S. (2020). Signaling models for dopamine-dependent temporal contiguity in striatal synaptic plasticity. PLOS Computational Biology, 16(7), e1008078. https://doi.org/10.1371/journal.pcbi.1008078

van Rijn, H. (2018). Towards Ecologically Valid Interval Timing. Trends in Cognitive Sciences, 22(10), 850–852. https://doi.org/10.1016/j.tics.2018.07.008

Wagenmakers, E.-J., Verhagen, J., & Ly, A. (2016). How to quantify the evidence for the absence of a correlation. Behavior Research Methods, 48(2), 413–426. https://doi.org/10.3758/s13428-015-0593-0

Wang, J., Hosseini, E., Meirhaeghe, N., Akkad, A., & Jazayeri, M. (2020). Reinforcement regulates timing variability in thalamus. ELife, 9. https://doi.org/10.7554/eLife.55872

Wiecki, T. V., Sofer, I., & Frank, M. J. (2013). HDDM: Hierarchical Bayesian estimation of the Drift-Diffusion Model in Python. Frontiers in Neuroinformatics, 7, 1–10. https://doi.org/10.3389/fninf.2013.00014

Wiener, M., Lee, Y.-S., Lohoff, F. W., & Coslett, H. B. (2014). Individual differences in the morphometry and activation of time perception networks are influenced by dopamine genotype. NeuroImage, 89, 10–22. https://doi.org/10.1016/j.neuroimage.2013.11.019

Wiener, M., Lohoff, F. W., & Coslett, H. B. (2011). Double Dissociation of Dopamine Genes and Timing in Humans. Journal of Cognitive Neuroscience, 23(10), 2811–2821. https://doi.org/10.1162/jocn.2011.21626

Zhou, S., Masmanidis, S. C., & Buonomano, D. V. (2020). Neural Sequences as an Optimal Dynamical Regime for the Readout of Time. Neuron, 108(4), 651-658.e5. https://doi.org/10.1016/j.neuron.2020.08.020

