## Supplementary material for "A proxy measure of striatal dopamine predicts individual differences in temporal precision": supplemental_mat.docx

Address correspondence to:

Renata Sadibolova

#### S1. Experimental task

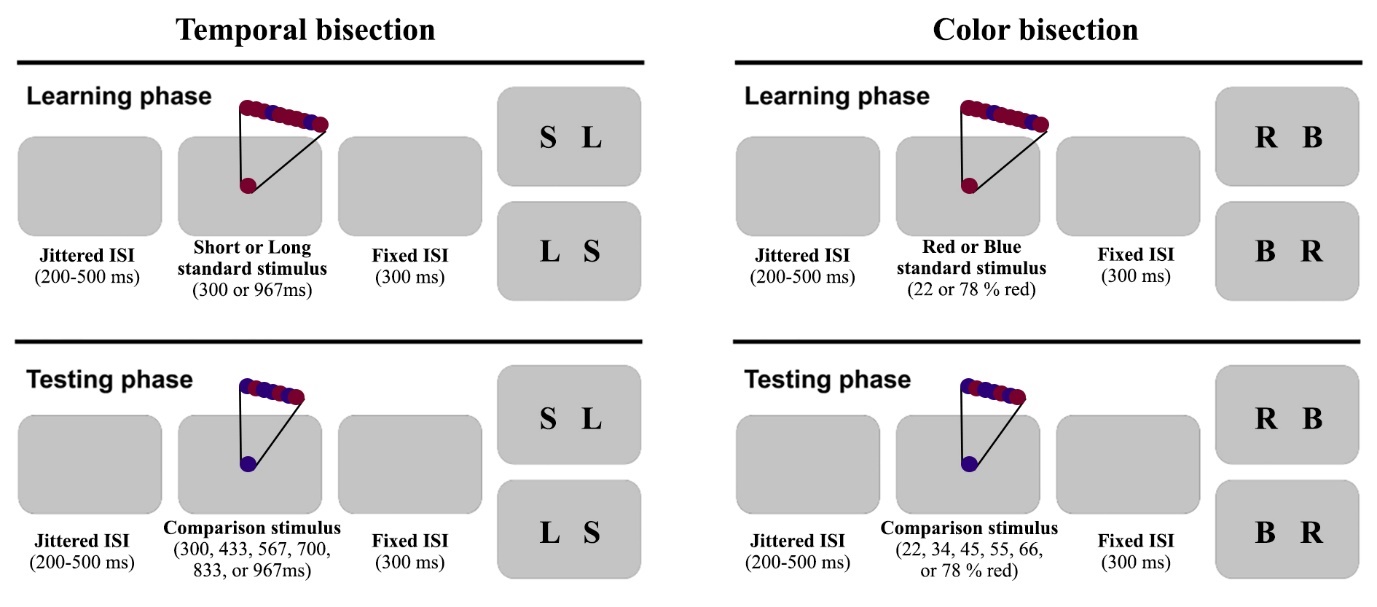

*Figure S1*. Experimental tasks. Participants observed a circle flickering between maroon and indigo colors at 60 Hz. Their task was to focus either on the duration (*Temporal bisection*) or the color (*Color bisection*). Each task comprised learning of a pair of standard stimuli followed by a testing phase in which participants judged the comparison stimuli with reference to the learnt standards. In the temporal bisection task, in the training phase, participants first learnt the short and long standard intervals (300 and 967ms). Subsequently, in the testing phase, they judged whether a comparison stimulus of varying duration (300, 433, 567, 700, 833, or 967ms) was closer in duration to the short or long standard interval. Each stimulus interval included an equal number of trials for color proportions investigated in the color bisection task. In the color bisection task, in the training phase participants learnt the relative proportion of the redder (vs. bluer) standards (22 and 78% red). In the testing phase, they were presented with circles of varying color proportion (22, 34, 45, 55, 66, or 78% red) and judged whether they were closer to the red or blue standard stimuli. Each color proportion included an equal number of trials for the intervals investigated in temporal bisection. All trials consistent of a blank screen (ISI of 200-500ms), a stimulus, and a post-stimulus empty screen (ISI of 300ms) followed by the response key mappings counterbalanced across participants. Participants responded by pressing the left and right arrows on a computer keyboard using their index and middle fingers, respectively.

#### S2. Bayes factor robustness

BFs are affected by the width of the H_1_ distribution prior (Wagenmakers et al., 2016). To assess the robustness of reported BFs based on a default JASP width *k*=1 (JASP Team, 2019), we compared how BF changes with varying prior distributions. When the width of a prior distribution equals zero, the H_1_=H_0_ and the BF=1. Prior belief that larger absolute correlation coefficient values are plausible under the H_1_ hypothesis is reflected in a wider prior; until all coefficients in the range of -1 to 0 (BF_-0_), 0 to 1 (BF_+0_), or -1 to 1 (BF_10_) are considered as equally probable at width *k*=2. Consequently, qualitatively similar BFs for different H_1_ priors attest to the reliability of the reported BF.

Our data showed reliably moderate evidence for EBR correlations with temporal bisection indices (Figure S2, top row). Figure S2 (top left) shows moderate evidence in support of H_1_ for a correlation of EBR and temporal WF (see main text). The robustness of Bayesian evidence would decrease for the prior width *k*>1.7, which incorporates correlation coefficients with large values close to 1. The top-right panel shows moderate Bayesian evidence in support of H_0_ for a correlation of EBR and temporal BP which is yet stronger for a larger prior width *k*>1.7. The evidence supporting H_0_ for EBR correlations with the color bisection indices was less robust and held only for prior width *k*>1 (bottom row). This suggests that H_0_ may hold only when large correlation coefficients are considered, and that the study might not be sufficiently powered to reliably detect evidence for the null hypothesis with smaller correlations in the color bisection task. Nevertheless, please refer to supplemental materials (section S6), where we report moderate evidence in favor of these null hypotheses after the exclusion of multivariate outliers.

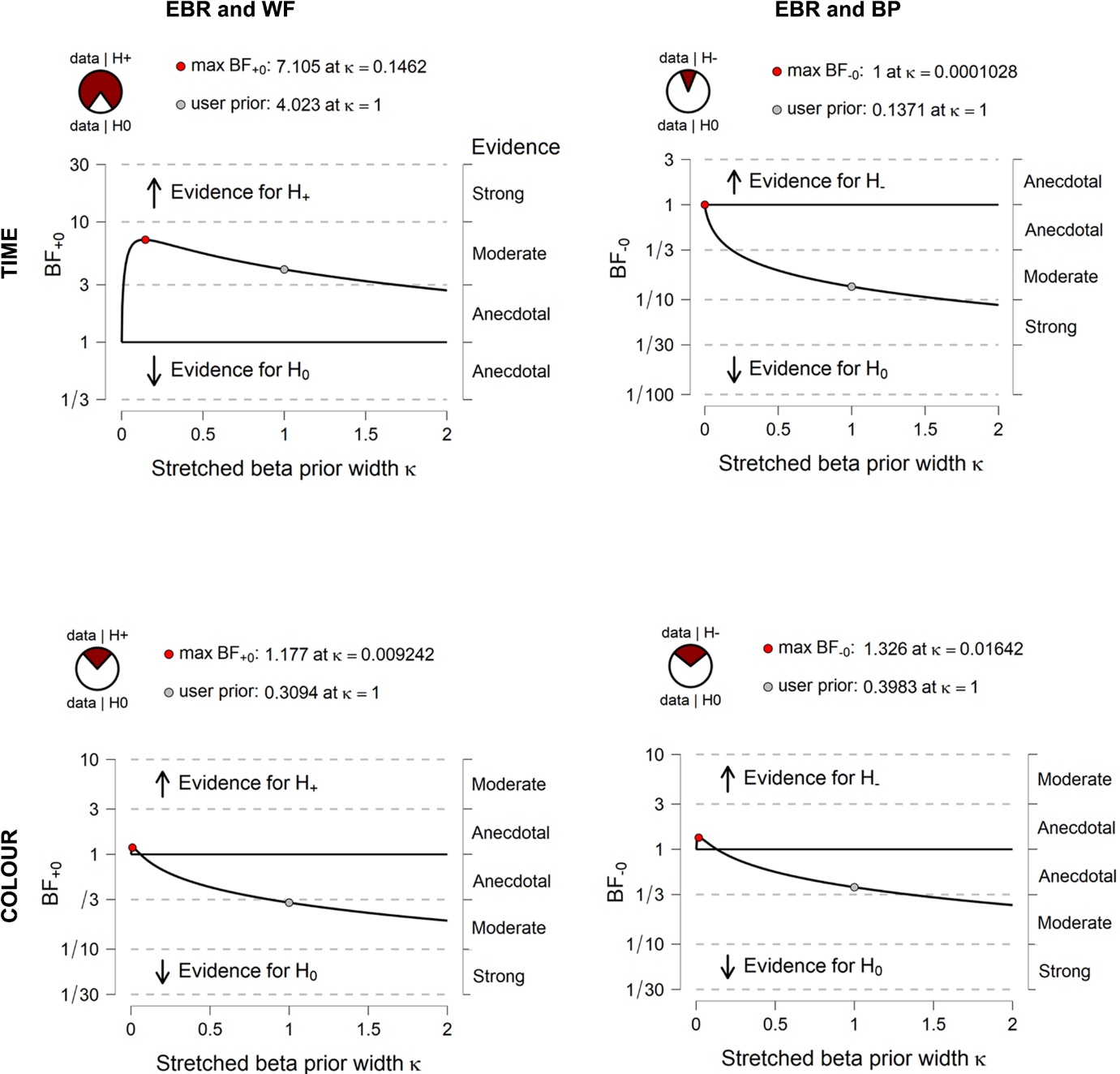

*Figure S2*. Bayes factor sensitivity analysis. BFs (*y axes*) were computed with H_1_ prior distributions of different widths (*x axes*) for the EBR/WF and EBR/BP correlations (left and right panels, respectively) in each task (panel rows). The BF_+0_ and BF_-0_ correspond to the hypothesized positive and negative H_1_ correlations, respectively. Reported BFs assume a default prior width of 1 (grey circles and pie charts). By quantifying the BF change as a function of different prior widths, the sensitivity analysis allows for robustness checks of BFs.

#### S3. Correlations of task precision and bias

In order to clarify patterns in our data and evaluate potential confounds, we conducted a series of exploratory correlation analyses. The WF and BP in temporal bisection showed negative correlation in line with the reward-rate maximization account (Balci et al., 2011), *r*_s_=-.24 [-.48, .01], *p*=.04 (*N*=69), BF_10_=1.19, and *r*_s_=-.30 [-.51, -.06], *p*=.02 (*N*=62), BF_10_=1.90 after the removal of bivariate outliers. However, it was supported only by anecdotal Bayesian evidence. Similarly, the strength of the evidence for negative correlation between the WF and BP in color bisection, *r*_s_=-.31, [-.53, -.09], *p*=.03 (*N*=67), BF_10_=4.10 was substantially reduced after the exclusion of bivariate outliers, *r*_s_=-.24 [-.46, .01], *p*=.06 (*N*=64), BF_10_=0.94. The evidence remained only anecdotal for a range of prior widths once the outliers were removed, with the maximum BF_10_=1.92 observed for k=0.08.

In order to assess that the former effect in the temporal bisection task did not confound the observed correlation between EBRs and temporal bisection WFs, we repeated this analysis adjusting for BPs and found that the correlation, *r*_ps_=.30 [.04, .49], *p*=.01 (*N*=69), BF_+0_=5.80, was supported by very strong Bayesian evidence after the exclusion of bivariate outliers, *r*_ps_=.41 [.19, .60], *p*<.001 (*N*=62), BF_+0_=51.58. By contrast, controlling for color bisection BPs did not improve the EBR and WF correlation in the color bisection task, *r*_ps_=.05 [-.19, .30], *p*=.68 (*N*=67), BF_+0_=.24, including without the outliers, *r*_ps_=.03 [-.25, .25], *p*=.84 (*N*=64), BF_+0_=.20.

Finally, EBR and BP correlations remained non-significant in both tasks after controlling for WF, *r*_ps_=.08 [-.18, .34], *p*=.51 (*N*=69), BF_10_=.18 (temporal bisection) and *r*_ps_=-.10 [-.35, .13], *p*=.41 (*N*=67), BF_10_=.22 (color bisection). These results remained stable after exclusion of bivariate outliers, *r*_ps_=.05 [-.21, .31], *p*=.70 (*N*=65), BF_10_=17 (temporal bisection) and *r*_ps_=-.07 [-.30, .15], *p*=.59 (*N*=61), BF_10_=19 (color bisection). Collectively, these results suggest that whilst temporal bisection WFs were associated with BPs, the observed association between WFs and EBR remained significant even when partialling out temporal bisection BPs whereas other exploratory analyses largely corroborated the other non-significant results in the main paper.

#### S4. Diffusion modelling

To further elucidate aspects of performance that may play a role in the reported effects, we fitted a drift diffusion model (DDM) (Ratcliff & McKoon, 2008; Ratcliff & Rouder, 1998) to the responses and response times in each task. The DDM models two-choice decision making as a noisy process of evidence accumulation over time. The two choices represent the upper and lower boundaries of a model, one of which is crossed by the evidence accumulation curve that drifts from the starting point adjusted by a bias parameter (*z*). The process approaches the boundary at a certain speed (drift rate; *v*), influenced by the amount of available evidence and noise that account for variable inter-trial boundary-crossings and response times. The distance between the boundaries (decision threshold; *a*) dictates how much evidence must be accumulated; with smaller distances leading to faster responding. A non-decision time parameter (*t*) represents ancillary non-decisional processes such as motor initiation and execution.

Our objective was to assess the relationship between EBR and different parameters of the DDM reflecting distinct elements of the decision process in each task. DDM parameters were estimated for each participant and at the group level using Bayesian inference and Markov Chain Monte Carlo (MCMC) sampling in the hierarchical drift diffusion modelling (HDDM) toolbox implemented in Python (Wiecki et al., 2013). We deviated from a preregistered temporal DDM model for two important reasons. First, it was desirable to apply an identical model fitting procedure in our non-temporal task, which could not be done with the temporal DDM model. Second, despite the advantages of the DDM tailored to temporal perception, the evidence shows that the standard DDM describes well the performance in the temporal bisection task that was used in our experiment (Balci & Simen, 2014, 2016; Wiener et al., 2018). Hierarchical models were compared by means of a deviance information criterion (DIC; Spiegelhalter et al., 2002) with its decrease by more than 10 accepted as sufficient evidence for model improvement.

In each task, we compared an “empty” model in which the parameters varied only by participant to the model including the stimulus (6 levels) as a continuous variable. This step was taken to establish that the latter model provided a better fit to the data. Each model was built using a chain of 10,000 samples from the posterior distribution, with the first 1,000 samples discarded and only every 5th sample retained thereafter to reduce sample autocorrelation and increase chain stability. The chains of each model were inspected to confirm symmetrical traces and distributions and low sample autocorrelations. A formal test of model convergence, the Gelman-Rubin statistic (GR; Gelman & Rubin, 1992) was computed with five different runs of each model, each drawing 5,000 samples with the first 200 discarded. The convergence issues are indicated by output values of >1.2.Following these procedures, we obtained the model parameter estimates (*a, v, t* and *z*) for each task which were correlated with EBRs using the robust correlation toolbox in MATLAB (Pernet et al., 2013).

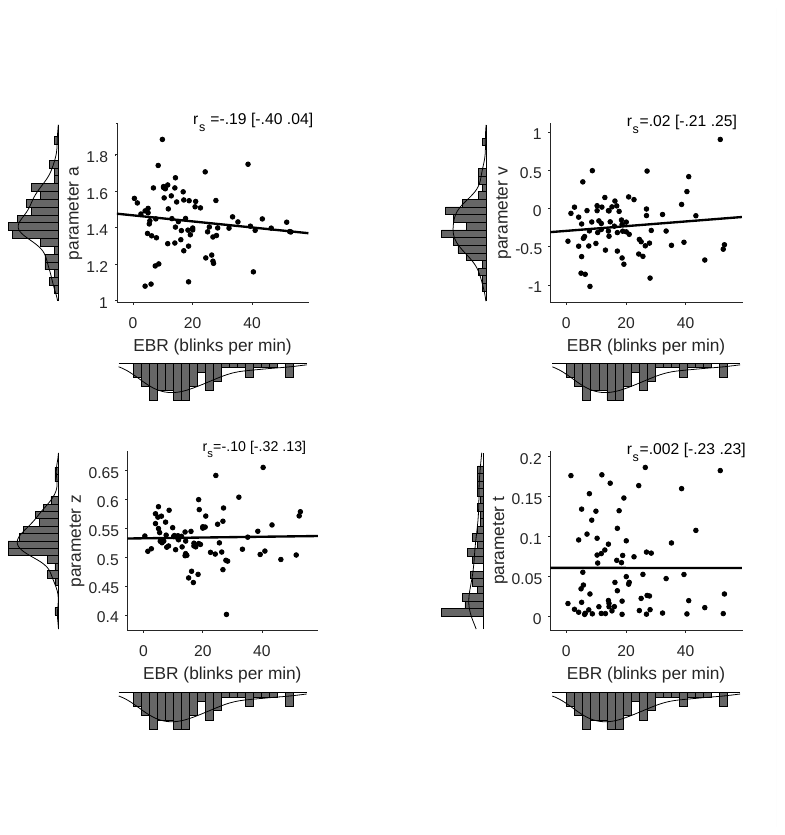

*Figure S3*: Scatterplots depicting the association between resting state spontaneous eyeblink rate (EBR) and temporal bisection task model parameters derived by a HDDM (*a*=decision threshold, *v*=drift rate, *z*=bias, *t*=non-decision time parameter). Square brackets denote Bootstrapped 95% confidence intervals. The scatter plots show non-ranked data with the least-squares line fit for visualization purposes.

The analysis showed that a better fitting model included the stimulus levels (DIC=15,991.55) relative to an empty model (24,103.75). Both models converged well according to symmetrical traces and minimal autocorrelations (the plots for these effects can be found at <https://osf.io/jxc3f/>). GR test values further corroborated the model convergence (M: 1.001, SD: .006). Individual GR test values were below the 1.02 cut-off except for estimates of the *t* parameter in five participants and for the *a* parameter in two participants. The analyses revealed that EBR did not significantly correlate with any of the HDDM parameters, including after the removal of bivariate outliers (see Table S1).

Table S1

Correlation coefficients between EBR and HDDM model parameters in the temporal and color bisection tasks.

| HDDM parameters | *r*_s_ [95% CI] | *p* | *N* | BF_(01 or +1)_ |
| --- | --- | --- | --- | --- |
| Temporal bisection |  |  |  |  |
| *a* | -.19 [-.40, .04] | .10 | 73 | .61 |
| *a* _EBO_ | -.17 [-.40, .06] | .16 | 69 | .43 |
| *v* | .02 [-.21, .25] | .86 | 73 | **.15** |
| *v* _EBO_ | .07 [-.18, .32] | .55 | 69 | **.18** |
| *z* | -.10 [-.32, .13] | .39 | 73 | **.26** |
| *z* _EBO_ | -.15 [-.40, .10] | .20 | 70 | .41 |
| *t* | .002 [-.23, .23] | .99 | 73 | **.15** |
| *t* _EBO_ | .01 [-.25, .25] | .96 | 70 | **.17** |
| Color bisection |  |  |  |  |
| *a* | -.05 [-.28, .18] | .66 | 73 | **.17** |
| *a* _EBO_ | -.03 [-.24, .19] | .84 | 70 | **.16** |
| *v* | .15 [-.08, .37] | .20 | 73 | **.27** |
| *v* _EBO_ | .12 [-.11, .34] | .33 | 70 | **.21** |
| *z* | -.15 [-.37, .08] | .21 | 73 | .38 |
| *z* _EBO_ | -.18 [-.39, .05] | .14 | 70 | .55 |
| *t* | .07 [-.17, .29] | .58 | 73 | **.17** |
| *t* _EBO_ | -.07 [-.15, .29] | .55 | 71 | **.18** |

*Notes.*	Bold values denote BF values reflecting moderate evidence for the null hypothesis.

EBR = eyeblink rate, HDDM = hierarchical drift diffusion model,

*a* = decision threshold, *v* = drift rate, *z* = bias, *t* = non-decision time,

EBO = excluding bivariate outliers.

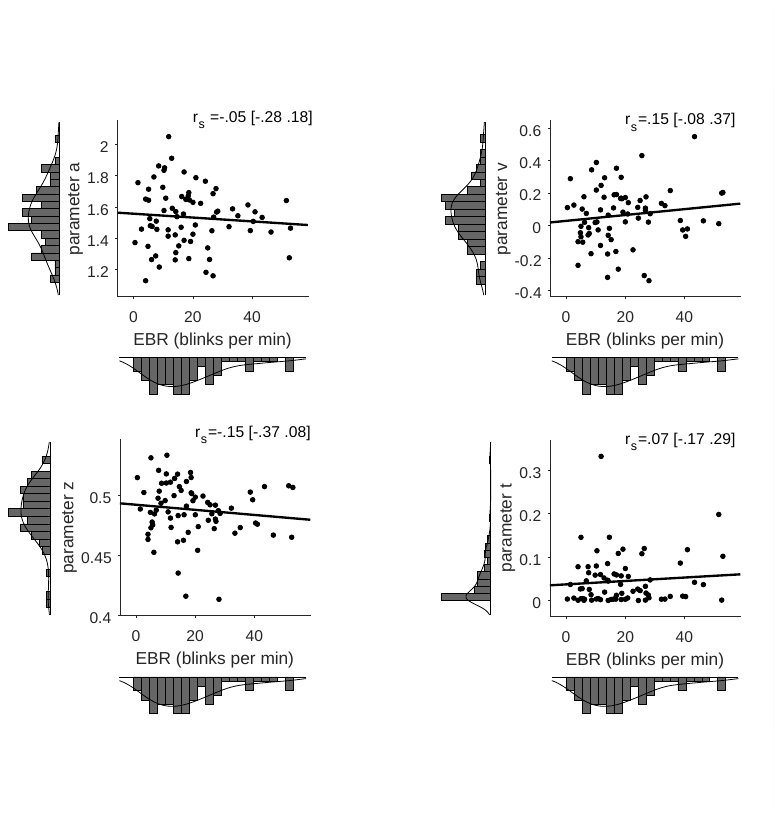

*Figure S4:* Scatterplots depicting the association between resting state spontaneous eyeblink rate (EBR) and color bisection task model parameters derived by a HDDM (*a*=decision threshold, *v*=drift rate, *z*=bias, *t*=non-decision time parameter). Square brackets denote Bootstrapped 95% confidence intervals. The scatter plots show non-ranked data with the least-squares line fit for visualization purposes.

The model including the color proportions (DIC = 20,025.00) better fitted the data in the color bisection task compared to the empty model (DIC = 28,961.72). Both models converged well according to symmetrical traces and minimal autocorrelations (for plots, see <https://osf.io/jxc3f>). Good convergence was further confirmed with the GR test values falling below the cut-off value of 1.02 (M: 1.002, SD: .006). Individual GR values above the cut-off include those for three *t* parameter estimates (red color proportions of 22, 34 and 45%) and nine participants. Our analyses revealed that EBR did not significantly correlate with HDDM parameters in the color bisection task (see Table S1).

Our final set of analyses comprised four paired-samples *t*-tests and correlations for each HDDM parameter pair in order to assess how the tasks differ in terms of the underlying decisional processes. The starting point for decision processes differed across tasks with a larger bias (*z*) in the temporal bisection task, *t*(74)=9.33, *p*<.001, *d*_z_=1.08. The *z* parameters across tasks did not correlate, *r*_ps_=.02 [-.22, .25], *p*=.88 (*N*=75), BF_10_=.34. Relative to the color bisection task, the evidence for timing decisions accumulated towards lower thresholds (*a*), *t*(74)=5.50, *p*<.001, *d*_z_=.64, and at a slower rate (*v*), *t*(74)=6.60, *p*<.001, *d*_z_=.76. However, whereas the drift rate correlation was non-significant, *r*_ps_=.06 [-.15, .30], *p*=.59 (*N*=75), BF_10_=.24, the thresholds across tasks did correlate, *r*_ps_=.61 [.42, .76], *p*<.001 (*N*=75), BF_10_=2.102e^+6^. Finally, the non-decision time parameter (t), representing ancillary non-decisional processes such as motor initiation and execution, was higher in the temporal bisection task, *t*(74)=2.67, *p*<.009, *d*_z_=.31 and correlated across tasks, *r*_ps_=.65 [.48, .77], *p*<.001 (*N*=75), BF_10_=1.263e^+6^. Altogether, these results suggest divergent decisional processes across experimental tasks.

#### S5. EBR and ambiguous trials

In previous research, low dopamine levels in Parkinson’s disease were associated with faster RTs for ambiguous (difficult) stimulus intervals (i.e., those closer to the BP), which was attributed to increased impulsive decision making (Zhang et al., 2016). To assess whether low dopamine activity, as inferred from low EBRs, was associated with faster RTs on ambiguous temporal bisection trials in our sample of neurotypical participants (Figure S.5), we correlated the differences of the largest and smallest mean stimulus level RT and EBRs. EBRs did not correlate with RT differences, *r*_s_=-.02 [-.25, .20], *p*=.89 (*N*= 73), BF_10_=.16 (no bivariate outliers). The analogous correlation in the color bisection task also did not reach significance, *r*_s_=-.02 [-.25, .21], *p*=.90 (*N*= 73), BF_10_=.16, including after removal of bivariate outliers, *r*_s_=.01 [-.22, .25], *p*=.91 (*N*=70), BF_10_=.18. These results suggests that unlike in Parkinson’s disease (Zhang et al., 2016), striatal dopamine receptor availability in healthy individuals, as inferred from EBR, is not associated with more impulsive decisions, with moderate evidence in favor of the null hypothesis.

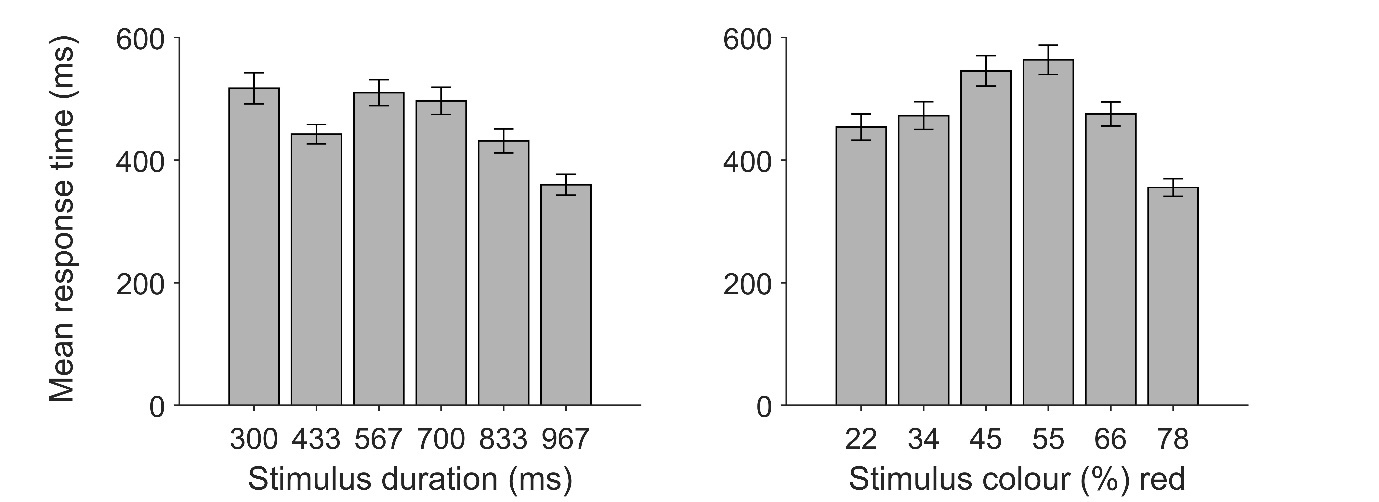

*Figure S5:* Mean response time for each stimulus level in temporal bisection task (left panel) and color bisection task (right panel).

#### S6. Bivariate outliers

Outlier detection was implemented using the ‘boxplot’ option of the robust correlation toolbox (Pernet et al., 2013). The algorithm detects outliers automatically by computing the distances to the center of a bivariate cloud using a projection method and subsequently eliminating values above the boxplot distance threshold (Wilcox, 2012).

The exclusion of four bivariate outliers strengthened the observed association between EBRs and WFs in the temporal bisection task, *r*_s_=.31 [.06, .53], *p*=.011 (*N*=65), BF_+0_=6.88. Similarly, we observed stronger Bayesian evidence in favor of the null hypothesis of no association between EBR and WF in the control (color bisection) task after excluding three bivariate outliers, *r*_s_=.04 [-.21, .29], *p*=.74 (N=64), BF_+0_=.22. Our results suggest that although temporal and color precision were associated, elevated striatal dopamine availability is selectively associated with poorer temporal precision. Moreover, the exclusion of nine bivariate outliers, *r*_s_=.28 [.02, .50], *p*=.036 (*N*=56), BF_+0_=3.00, did not affect the across-task correlation between WFs. Importantly, a semi-partial Spearman correlation between temporal bisection WFs and EBR (after partialling out shared variance between the temporal and color bisection WFs), remained significant including after removal of the bivariate outliers, *r*_ps_ =.31 [.04, .53], *p*=.018 (*N*=58), BF_+0_=5.77. Conversely, the non-significant correlation between EBR and color bisection WFs did not change after controlling for temporal bisection WFs, including without the bivariate outliers, *r*_ps_ =-.11 [-.35, .14], *p*=.394 (*N*=61), BF_+0_=.10.

In contrast with our second prediction regarding perceived duration, our results suggest that inter-individual differences in baseline striatal dopamine availability, as inferred from EBR, are unrelated to inter-individual variation in perceived duration. EBR did not significantly correlate with temporal BPs, including after removal of bivariate outliers, *r*_s_=-.004 [-.26, .25], *p*=.97 (*N*=65), BF_-0_=.18. Similarly, the removal of six bivariate outliers in the color bisection task did not result in a significant EBR and BP correlation, *r*_s_=.04, [-.22, .28], *p*=.77 (*N*=61), BF_-0_=.12. Finally, BPs in the two tasks did not significantly correlate after the removal of bivariate outliers, *r*_s_=.15, [-.13, .40], *p*=.24 (*N*=63), BF_10_=.33.

#### S7. The strength of association between EBR and dopamine D2-receptors

Eyeblinks are readily observed and EBRs are easily quantified. Their use as a proxy striatal dopamine measure circumvents multiple challenges imposed by alternative methods that are either invasive or include pharmacological interventions with temporo-spatially diffuse effects in the brain. It is nevertheless critical to acknowledge the shortcomings of this indirect measure which fundamentally depends on the strength of the association between EBR and striatal D2-receptor availability. Whereas the prolific literature bearing on this association has been comprehensively reviewed elsewhere (Jongkees & Colzato, 2016), we summarize in Table S2 the studies that recorded EBR and dopamine or D2 receptor function directly using PET so as to provide additional details regarding the strength of their correlation.

Table S2

The association of EBR and dopamine/D2-receptor function.

| Publication | Findings |
| --- | --- |
| * Dang et al. (2017) | *N*=20. No correlation between EBR and **D2-receptor availability (PET)** in caudate (β= –0.21), putamen (β= –0.22), ventral striatum (β= 0.24), and midbrain (β= 0.04). |
| Groman et al. (2014) | *N*=10. The association between resting EBR and **D2-receptor density (PET and postmortem)** in caudate nucleus (*r*=0.65), putamen (*r*=0.58) and ventral striatum (*r*=0.74) which was stable within and across assessments 3 months apart (within-session reliability, Cronbach’s alpha >0.87; between-session correlation *r*=0.81 ). |
| Sescousse et al. (2018) | *N*=20. Negative association between EBR and **dopamine synthesis capacity (PET)** (Spearman *r*=-0.50) in left nucleus accumbens. |
| Taylor et al. (1999) | *N*=9. Positive association between EBR and **dopamine concentration (postmortem)** in ventromedial caudate striatum (*r*=.80). |
| Verhoeff et al. (2003) | *N*=6. Positive correlation for post-amphetamine EBR and striatal **D2-receptor binding potential (PET)** (*r*=.85). |

*Notes.* ***** PET data were acquired **17 [range: 3 - 32] months** before EBR (Dang et al., 2017).

### References

Balci, F., Freestone, D., Simen, P., Desouza, L., Cohen, J. D., & Holmes, P. (2011). Optimal temporal risk assessment. *Frontiers in Integrative Neuroscience*, *5*. https://doi.org/10.3389/fnint.2011.00056

Balci, F., & Simen, P. (2014). Decision processes in temporal discrimination. *Acta Psychologica*, *149*, 157–168. https://doi.org/10.1016/j.actpsy.2014.03.005

Balci, F., & Simen, P. (2016). A decision model of timing. *Current Opinion in Behavioral Sciences*, *8*, 94–101. https://doi.org/10.1016/j.cobeha.2016.02.002

Dang, L. C., Samanez-Larkin, G. R., Castrellon, J. J., Perkins, S. F., Cowan, R. L., Newhouse, P. A., & Zald, D. H. (2017). Spontaneous Eye Blink Rate (EBR) Is Uncorrelated with Dopamine D2 Receptor Availability and Unmodulated by Dopamine Agonism in Healthy Adults. *Eneuro*, *4*(5), ENEURO.0211-17.2017. https://doi.org/10.1523/ENEURO.0211-17.2017

Gelman, A., & Rubin, D. B. (1992). Inference from Iterative Simulation Using Multiple Sequences. *Statistical Science*, *7*(4), 457–472. https://doi.org/10.1214/ss/1177011136

Groman, S. M., James, A. S., Seu, E., Tran, S., Clark, T. A., Harpster, S. N., Crawford, M., Burtner, J. L., Feiler, K., Roth, R. H., Elsworth, J. D., London, E. D., & Jentsch, J. D. (2014). In the Blink of an Eye: Relating Positive-Feedback Sensitivity to Striatal Dopamine D2-Like Receptors through Blink Rate. *Journal of Neuroscience*, *34*(43), 14443–14454. https://doi.org/10.1523/JNEUROSCI.3037-14.2014

JASP Team. (2019). JASP. In *[Computer software]*.

Jongkees, B. J., & Colzato, L. S. (2016). Spontaneous eye blink rate as predictor of dopamine-related cognitive function—A review. *Neuroscience & Biobehavioral Reviews*, *71*, 58–82. https://doi.org/10.1016/j.neubiorev.2016.08.020

Pernet, C. R., Wilcox, R., & Rousselet, G. A. (2013). Robust Correlation Analyses: False Positive and Power Validation Using a New Open Source Matlab Toolbox. *Frontiers in Psychology*, *3*. https://doi.org/10.3389/fpsyg.2012.00606

Ratcliff, R., & McKoon, G. (2008). The Diffusion Decision Model: Theory and Data for Two-Choice Decision Tasks. *Neural Computation*, *20*(4), 873–922. https://doi.org/10.1162/neco.2008.12-06-420

Ratcliff, R., & Rouder, J. N. (1998). Modeling Response Times for Two-Choice Decisions. *Psychological Science*, *9*(5), 347–356. https://doi.org/10.1111/1467-9280.00067

Sescousse, G., Ligneul, R., van Holst, R. J., Janssen, L. K., de Boer, F., Janssen, M., Berry, A. S., Jagust, W. J., & Cools, R. (2018). Spontaneous eye blink rate and dopamine synthesis capacity: preliminary evidence for an absence of positive correlation. *European Journal of Neuroscience*, *47*(9), 1081–1086. https://doi.org/10.1111/ejn.13895

Spiegelhalter, D. J., Best, N. G., Carlin, B. P., & van der Linde, A. (2002). Bayesian measures of model complexity and fit. *Journal of the Royal Statistical Society: Series B (Statistical Methodology)*, *64*(4), 583–639. https://doi.org/10.1111/1467-9868.00353

Taylor, J. R., Elsworth, J. D., Lawrence, M. S., Sladek, J. R., Roth, R. H., & Redmond, D. E. (1999). Spontaneous Blink Rates Correlate with Dopamine Levels in the Caudate Nucleus of MPTP-Treated Monkeys. *Experimental Neurology*, *158*(1), 214–220. https://doi.org/10.1006/exnr.1999.7093

Verhoeff, N. P. L. ., Christensen, B. K., Hussey, D., Lee, M., Papatheodorou, G., Kopala, L., Rui, Q., Zipursky, R. B., & Kapur, S. (2003). Effects of catecholamine depletion on D2 receptor binding, mood, and attentiveness in humans: a replication study. *Pharmacology Biochemistry and Behavior*, *74*(2), 425–432. https://doi.org/10.1016/S0091-3057(02)01028-6

Wagenmakers, E.-J., Verhagen, J., & Ly, A. (2016). How to quantify the evidence for the absence of a correlation. *Behavior Research Methods*, *48*(2), 413–426. https://doi.org/10.3758/s13428-015-0593-0

Wiecki, T. V., Sofer, I., & Frank, M. J. (2013). HDDM: Hierarchical Bayesian estimation of the Drift-Diffusion Model in Python. *Frontiers in Neuroinformatics*, *7*, 1–10. https://doi.org/10.3389/fninf.2013.00014

Wiener, M., Parikh, A., Krakow, A., & Coslett, H. B. (2018). An intrinsic role of beta oscillations in memory for time estimation. *Scientific Reports*. https://doi.org/10.1038/s41598-018-26385-6

Wilcox, R. R. (2012). Introduction to Robust Estimation and Hypothesis Testing. In *Introduction to Robust Estimation and Hypothesis Testing* (3rd Ed.). Elsevier Inc. https://doi.org/10.1016/C2010-0-67044-1

Zhang, J., Nombela, C., Wolpe, N., Barker, R. A., & Rowe, J. B. (2016). Time on timing: Dissociating premature responding from interval sensitivity in Parkinson’s disease. *Movement Disorders*, *31*(8), 1163–1172. https://doi.org/10.1002/mds.26631
